# Respiratory syncytial virus infection confers heterologous protection against SARS-CoV-2 via induction of γδ T cell-mediated trained immunity and SARS-CoV-2 reactive mucosal T cells

**DOI:** 10.1101/2025.07.02.662833

**Authors:** Awadalkareem Adam, Wenzhe Wu, Madison C Jones, Haiping Hao, Aditi, Parimal Samir, Xiaoyong Bao, Tian Wang

**Affiliations:** Department of Microbiology & Immunology, University of Texas Medical Branch, Galveston, TX, 77555, USA; Sealy Institute for Vaccine Sciences, University of Texas Medical Branch, Galveston, TX 77555, USA; Department of Pediatrics, University of Texas Medical Branch, Galveston, TX 77555, USA; Department of Biochemistry & Molecular Biology, University of Texas Medical Branch, Galveston, TX 77555, USA; Institute for Human Infections and Immunity, University of Texas Medical Branch, Galveston, TX, 77555, USA; Department of Pathology, University of Texas Medical Branch, Galveston, TX, 77555, USA

**Keywords:** RSV, trained immunity, SARS-CoV-2, γδ T cells

## Abstract

The respiratory viruses can concurrently or sequentially infect a host and influence the trajectory of each other. The underlying immune mechanisms are not well understood. Here, we investigated whether respiratory syncytial virus (RSV) infection affects host vulnerability to subsequent SARS-CoV-2 infection in two murine models of SARS-CoV-2 infection. We found that prior RSV infection-induced heterologous protection against subsequent SARS-CoV-2 infection was dose and time dependent. RNA-seq and immunological analyses revealed that RSV triggered the activation of lung antigen presenting cells (APC)s and SARS-CoV-2 reactive mucosal T cells at day 9, which declined at 1 month. RSV also induced the expansion of lung γδ T cells and the upregulation of their cellular metabolic pathways. Furthermore, RSV infection in TCRδ^-/-^ mice, which are deficient of γδ T cells, resulted in a reduced SARS-CoV-2 reactive mucosal T cell response and subsequent increased viral loads and higher levels of virus-induced inflammatory responses in the lung upon SARS-CoV-2 challenge compared to the wild-type mice. In summary, our findings suggest that RSV infection provides heterologous protection against the subsequent SARS-CoV-2 infection via induction of γδ T cell-mediated trained immunity in the lung and SARS-CoV-2 reactive mucosal T cell responses.

## INTRODUCTION

Severe Acute Respiratory Syndrome Coronavirus 2 (SARS-CoV-2) was the cause of the recent coronavirus disease 2019 (COVID-19) pandemic, which had made a devastating impact on global public health for more than four years. The virus continues to evolve and circulate in human population. In addition to SARS-CoV-2, several other respiratory viral infections, such as influenza, respiratory syncytial virus (RSV), metapneumovirus (HMPV), rhinovirus (RV), are known to be the causative agents of acute respiratory diseases in infants, the elderly, and immunocompromised individuals. Clinical signs of these respiratory viral infections range from mild upper respiratory symptoms to more serious lower respiratory illnesses, including bronchiolitis and pneumonia. Additionally, these illnesses can have a long-lasting impact on patient health well beyond the resolution of the viral infection. These respiratory viruses could circulate with each other during winter and spring seasons and concurrently or sequentially infect a host and influence both host susceptibility and the severity of viral diseases by each other ^1^ via modulation host immunity. During the early COVID-19 pandemic, 20 to 50% of healthy donors with no prior exposure to SARS-CoV-2 were reported to have cross-reactive CD4^+^ T cells against SARS-CoV-2 and the common human coronaviruses (HCoV)s ^2^. Prior infection with RSV in mice was reported to be protected from the heterologous influenza virus-induced diseases via activation of alveolar macrophages, and RSV-specific CD8^+^ T cells ^3^. Whether prior infections with other respiratory viruses affect host vulnerability to SARS-CoV-2 disease and host immunity remains largely unknown.

Trained immunity is a long-term increase in responsiveness of innate immune cells via metabolic, epigenetic reprogramming, and transcriptional changes ^4,5^. Epidemiological studies over the years suggested that live attenuated vaccines, such as Bacillus Calmettte-Guerin (BCG) vaccine, Measles, mumps and rubella (MMR) vaccine, polio vaccine, and influenza vaccines confer nonspecific protection against unrelated infections via induction of trained immunity ^6–9^. Animal model studies have shown that cytomegalovirus or bacterial infections-induced trained immunity provides broad-spectrum protection against other microbial infections ^10–12^. In this study, we investigated prior RSV infection on host susceptibility against subsequent SARS-CoV-2 challenge in two animal models. Our findings suggest that RSV infection triggers γδ T cell-mediated trained immunity and promotes the induction of SARS-CoV-2-reactive mucosal T cells, which together provide heterogeneous protection against subsequent SARS-CoV-2 infection.

## RESULTS

### Prior RSV infection induced heterologous protection against subsequent SARS-CoV-2 infection is dose and time-dependent

To study the impacts of RSV infection on the subsequent SARS-CoV-2 challenge, we initially infected BALB/c mice intranasally (i.n.) with 5 x 10^6^ PFU of the RSV A2 strain or PBS (mock) and monitored them daily for weight loss and morbidity. Next, infected mice were i.n. challenged with 1 × 10^4^ PFU of the mouse-adapted SARS-CoV-2 strain CMA4 on day 9 post RSV infection (**Figure 1A**). The mouse-adapted strain infects the lungs, causes inflammatory responses, and reaches to infection peak around day 2 ^13,14^. To assess SARS-CoV-2-induced disease severity, we measured viral loads and chemokine levels in the lung 2 days after SARS-CoV-2 challenge. SARS-CoV-2-infected mice with prior RSV infection [SARS-CoV-2 (RSV)] showed markedly reduced viral loads in the lung as determined by Q-PCR and plaque assay (**Figure 1B-C**). There were 4 to 11-fold lower levels of chemokines, including *Ccl2*, *Cxcl10*, and *Ccl7* (**Figure 1D**) compared to SARS-CoV-2-infected mice with prior mock infection [SARS-CoV-2 (Mock)]. K18-hACE2 transgenic mice express the hACE2 protein under the human keratin 18 (K18) promoter and are in C57BL/6 (B6) genetic background, which confers efficient transgene expression in airway epithelial cells. Acute SARS-CoV-2 infection in K18-hACE2 transgenic mice induces more pronounced weight loss, interstitial pneumonitis, encephalitis, and death ^15–18^. To further investigate the impact of prior RSV infection on host susceptibility to subsequent SARS-CoV-2 infection, K18-hACE2 mice were next infected with a low dose (LD, 5 x10^6^ PFU) or high dose (HD, 1 x10^7^PFU) RSV. Infected mice were monitored daily for morbidity and weight changes.

**Figure 1.**
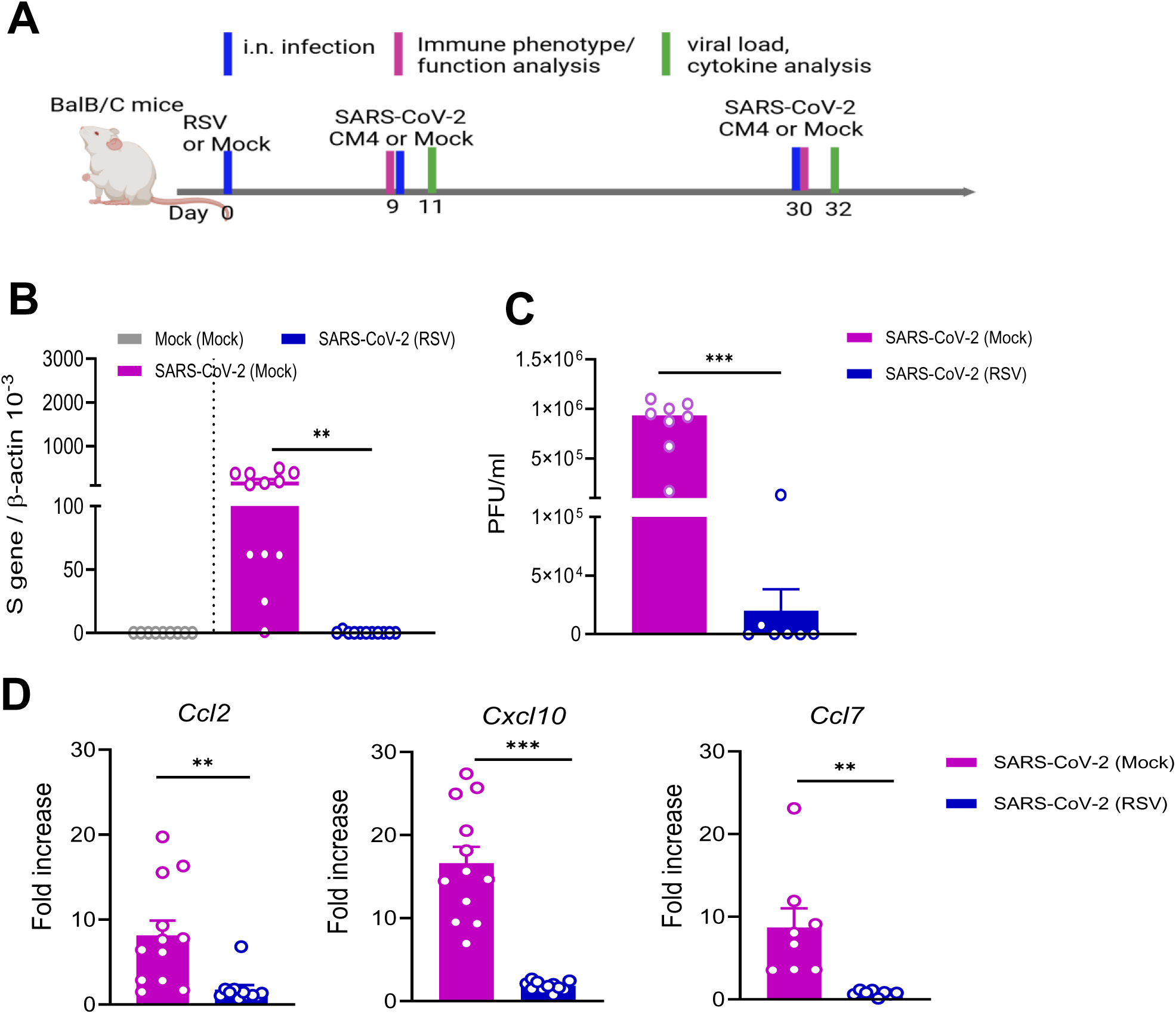
Prior RSV infection increased host resistance to subsequent SARS-CoV-2 infection in BALB/C mice. BALB/c mice were infected i.n. with RSV or PBS (mock), and on day 9, mice were challenged with mouse-adaptive SARS-CoV-2 strain CMA4. **A**. Study design (created with BioRender.com). **B-D.** Lung tissues were collected at day 2 post SARS-CoV-2 challenge. **B-C.** Lung viral load measured by Q-PCR of spike (S) gene expression (**B**) and plaque assays (**C**). **D**. Lung chemokine levels at day 2 post challenge determined by Q-PCR assay. Data are presented as fold increase compared to mock-infected mice. n = 9 to 12. \*\**P* < 0.01, or \*\*\**P* < 0.001 SARS-CoV-2 - infected mice with RSV prior infection [SARS-CoV-2 (RSV)] compared to SARS-CoV-2-infected mice with mock prior infection [SARS-CoV-2 (mock)].

Infection by both doses of RSV resulted in weight loss in K18-hACE2 mice within a week of infection (**Supplementary Figure 1**). On day 9 of the RSV infection, mice were challenged with 1 x10^3^ PFU SARS-CoV-2 prototype strain (**Figure 2A**). Mice with prior mock infection started to lose weight on day 6, and about 80% of them succumbed to SARS-CoV-2 infection within a 3-week interval. In contrast, none of the mice with prior LD or HD RSV infection displayed weight loss, and all survived subsequent SARS-CoV-2 infection (**Figure 2B-C**).

**Figure 2.**
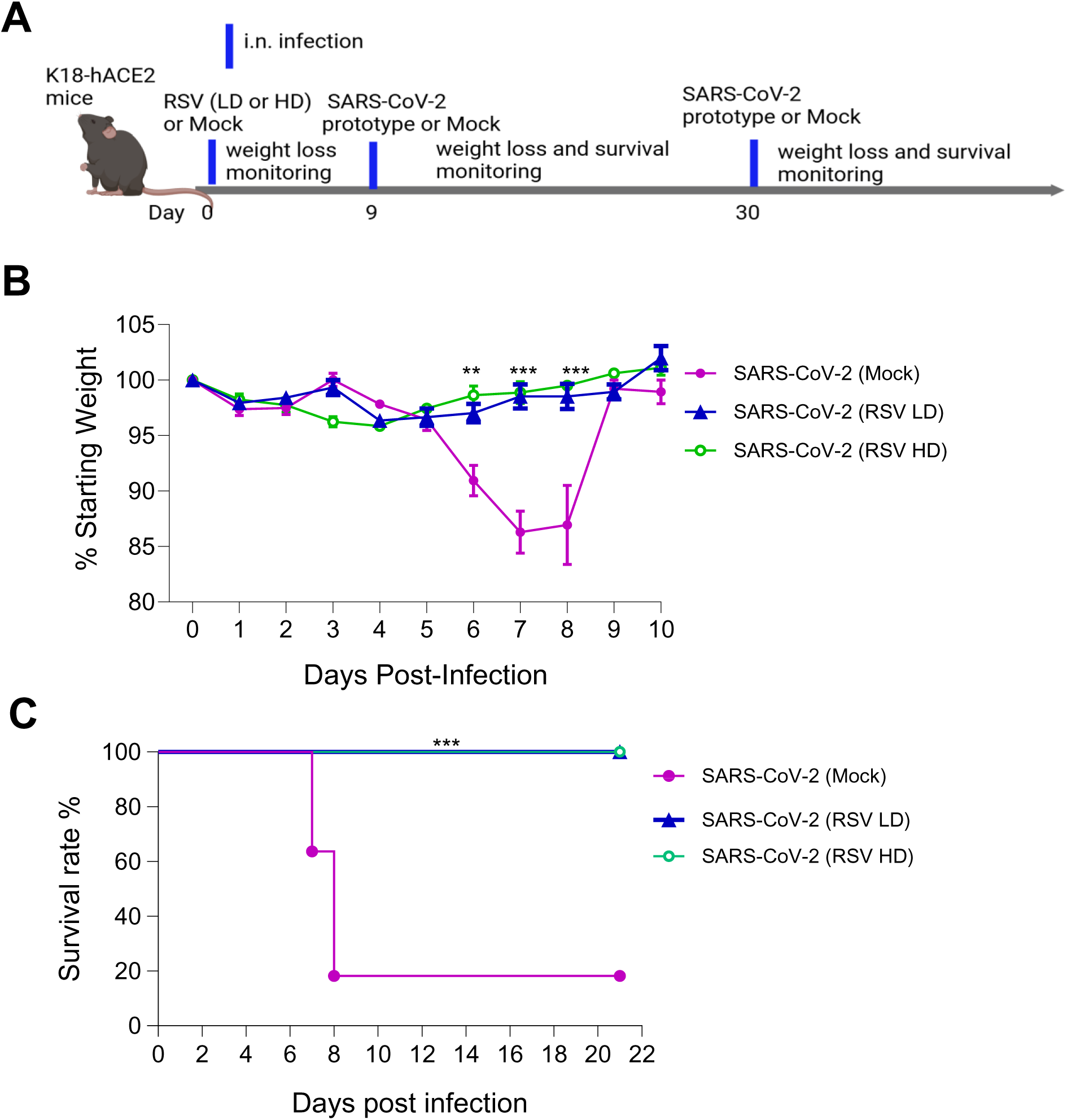
Prior RSV infection increased subsequent host survival from SARS-CoV-2 challenge in K18-hACE2 mice. K18-hACE2 mice were infected with a low (LD) or high dose (HD) of RSV or mock, and on day 9, mice were challenged with SARS-CoV-2 prototype strain. **A**. Study design (created with BioRender.com). Mice were monitored daily for weight loss (**B**) or survival (**C**). Weight loss is indicated by percentage using the weight on the day of infection as 100%. \*\*\**P* < 0.001 or \*\**P* < 0.01, [SARS-CoV-2 (RSV LD), n =13] or [SARS-CoV-2 (RSV HD), n= 6) compared to [SARS-CoV-2 (mock), n =8].

To determine the durability of RSV-induced heterologous protection, BALB/c mice were challenged with 1 x10^4^ PFU SARS-CoV-2 CMA4 30 days post RSV infection. At day 2 post SARS-CoV-2 challenge, viral loads in the lung were noted to be not significantly different between prior RSV-infected and prior mock-infected mice (**Figure 3A**). There were 1.5 to 3-fold lower levels of *Ccl7* and *Cxcl10* but similar levels of *Ccl2* in the lung of prior RSV infected mice than the mice with prior-mock-infection (**Figure 3B**). Next, K18-hACE2 mice were also challenged with 1 x10^3^ PFU SARS-CoV-2 prototype on day 30 post RSV (both LD and HD) or mock infection. All three groups displayed weight loss on day 6 of SARS-CoV-2 infection but significantly less weight loss in the RSV-prior infected LD and HD groups compared to those with mock prior infection (**Figure 3C**). On days 7 and 8, the HD RSV prior infected mice continued to display reduced weight loss compared to the mock prior infection group; while no significant differences in weight changes were noted between LD RSV-prior infection group and the mock prior infection group became insignificant. Furthermore, both LD (78.5% survival) and HD (85.7% survival) with prior RSV infection showed increased survival rates compared to mice prior infected with mock (30% survival, **Figure 3D**). Overall, these results suggest that prior RSV infection decreases host susceptibility to subsequent SARS-CoV-2 challenge in both mouse models. RSV-induced heterologous protection against subsequent SARS-CoV-2 challenge is dose and time-dependent.

**Figure 3.**
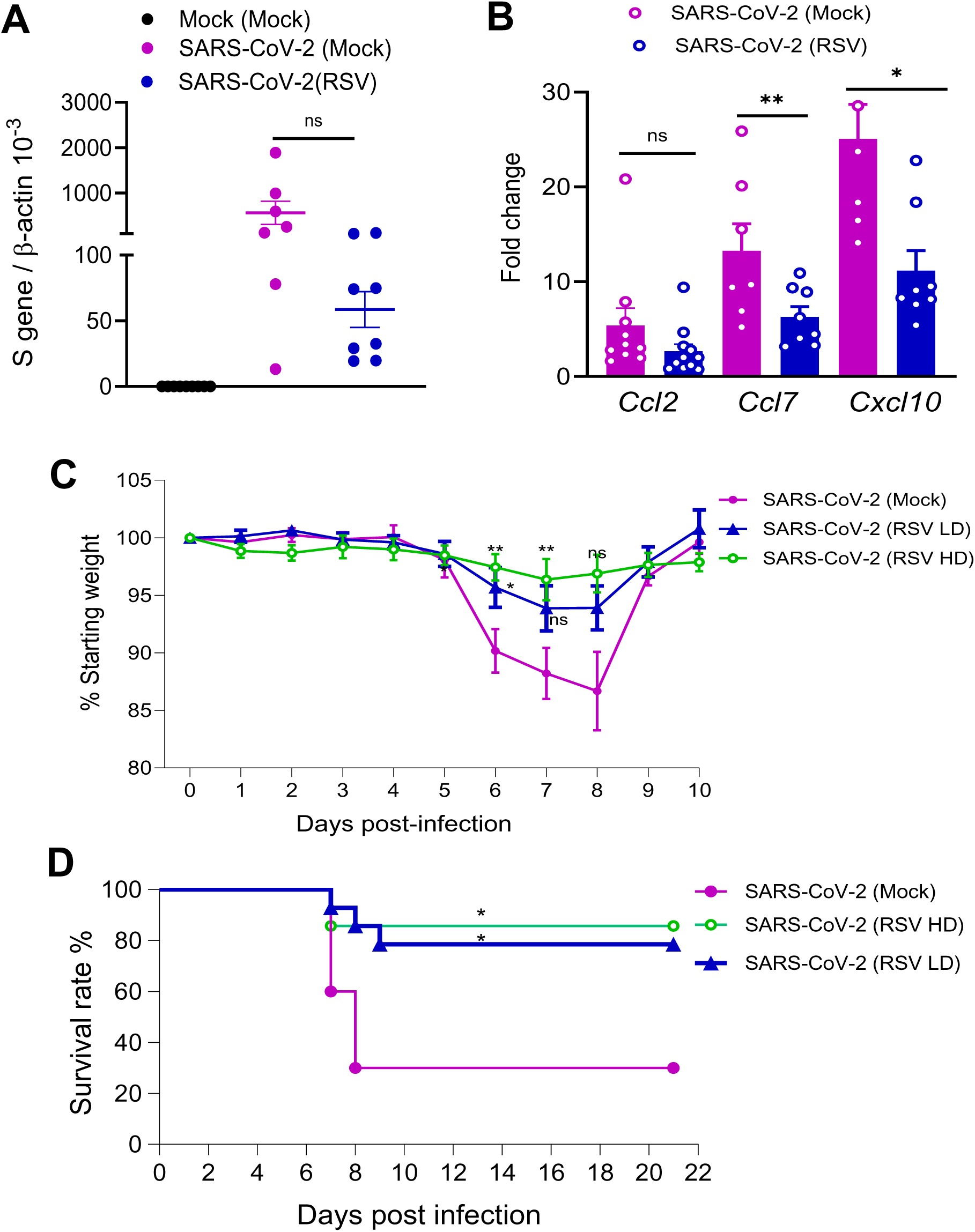
RSV-induced heterologous protection against SARS-CoV-2 declined at day 30 post RSV infection. **A-B.** BALB/c mice were infected i.n. with RSV, and on day 30 post infection, mice were challenged with mouse-adaptive SARS-CoV-2 strain CMA4. Lung tissues were collected at day 2 post SARS-CoV-2 challenge. Lung viral loads (**A**) and chemokine levels (**B**) were measured by Q-PCR assay. Data are presented as fold increase compared to mock-infected. n= 7 to 8. **C-D.** K18 hACE2 mice were infected with a low (LD) or high dose (HD) of RSV or mock, and on day 30 post RSV infection, mice were challenged with SARS-CoV-2 prototype strain. Mice were monitored daily for weight loss (**C**) and survival (**D**). Weight loss is indicated by percentage using the weight on the day of infection as 100%. \**P* < 0.05 or \*\**P* < 0.01 [SARS-CoV-2 (RSV LD), n = 14], [SARS-CoV-2 (RSV HD), n=6] compared to [SARS-CoV-2 (mock), n = 10)].

### Transcriptomic and functional analysis reveals that RSV infection triggered the activation of antigen presenting cells (APC) and SARS-CoV-2-reactive mucosal T cell responses

To understand the underlying immune mechanism of RSV-induced heterologous protection, we next infected B6 mice with RSV or mock, on day 9 post RSV infection, mice were challenged with 1 x10^4^ PFU SARS-CoV-2 CMA4 or mock. Lung tissue RNAs were isolated at day 9 post RSV infection and at day 2 post-SARS-CoV-2 challenge and subjected to transcriptomic analysis using RNA sequencing (seq). We found that RSV infection induced 100 top differentially expressed genes, were associated with multiple immune pathways such as “leukocyte cell-cell adhesion”, “regulation of T cell or lymphocyte activation”, “lymphocyte proliferation”, “regulation of innate immune responses”, “antigen receptor-mediated signaling pathways”, “type II interferon”, “innate Immunity activation”, “leukocyte activation” and “antigen processing” (**Figure 4A-B, and Supplementary Figure 2A**). Gene set enrichment analysis (GSEA) of the transcriptomics data set also revealed significant upregulation of immune-related pathways in the RSV-infected group compared to the mock group (**Supplementary Figure 2B-D**). Notably, pathways involved in “lymphocyte-mediated immunity”, “leukocyte-mediated immunity”, and “adaptive immune responses” showed marked enrichment in the RSV group. Specifically, the “lymphocyte-mediated immunity” pathway exhibited a strong positive enrichment score (NES =2.48), indicating elevated expression of genes associated with lymphocyte activation and effector functions in RSV-infected samples. Similarly, the “leukocyte-mediated immunity” pathway was significantly enriched (NES =2.40), suggesting enhanced recruitment and activation of leukocytes in response to RSV infection. The “adaptive immune response*”* pathway was also prominently upregulated (NES = 2.37), reflecting the activation of antigen-specific immune mechanisms, including T and B cell responses. These findings were further supported by corresponding heatmaps, which display higher expression levels of genes within these pathways in the RSV group compared to mock. Furthermore, similar sets of upregulated genes and associated induction of immune signaling pathways were noted when comparing SARS-CoV-2 infected mice with prior RSV infection to mock-infected mice (**Supplementary Figure 3**). Together, these data suggest that RSV-induced heterologous protection against SARS-CoV-2 is associated with activation of APCs and T cells.

**Figure 4.**
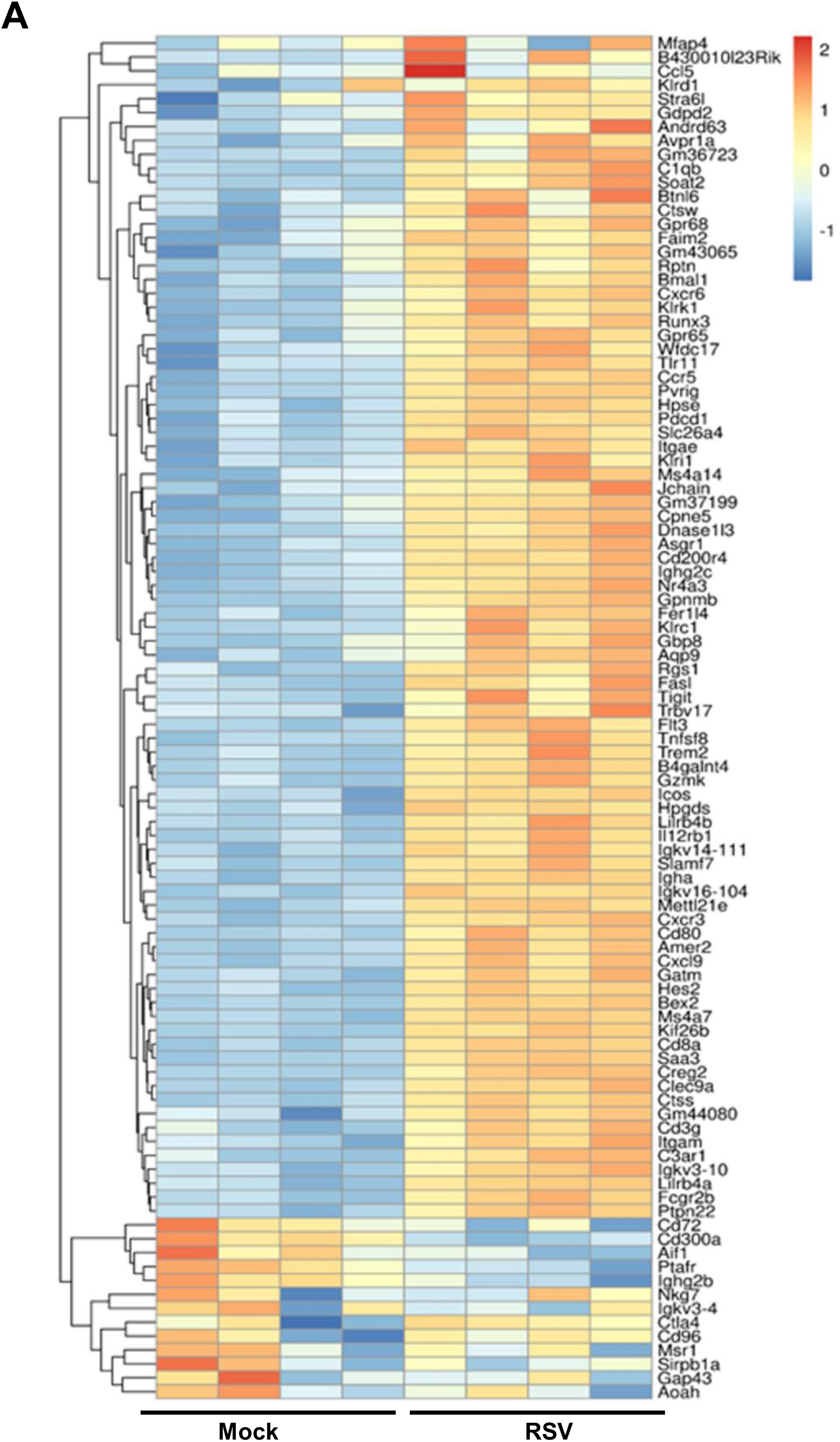
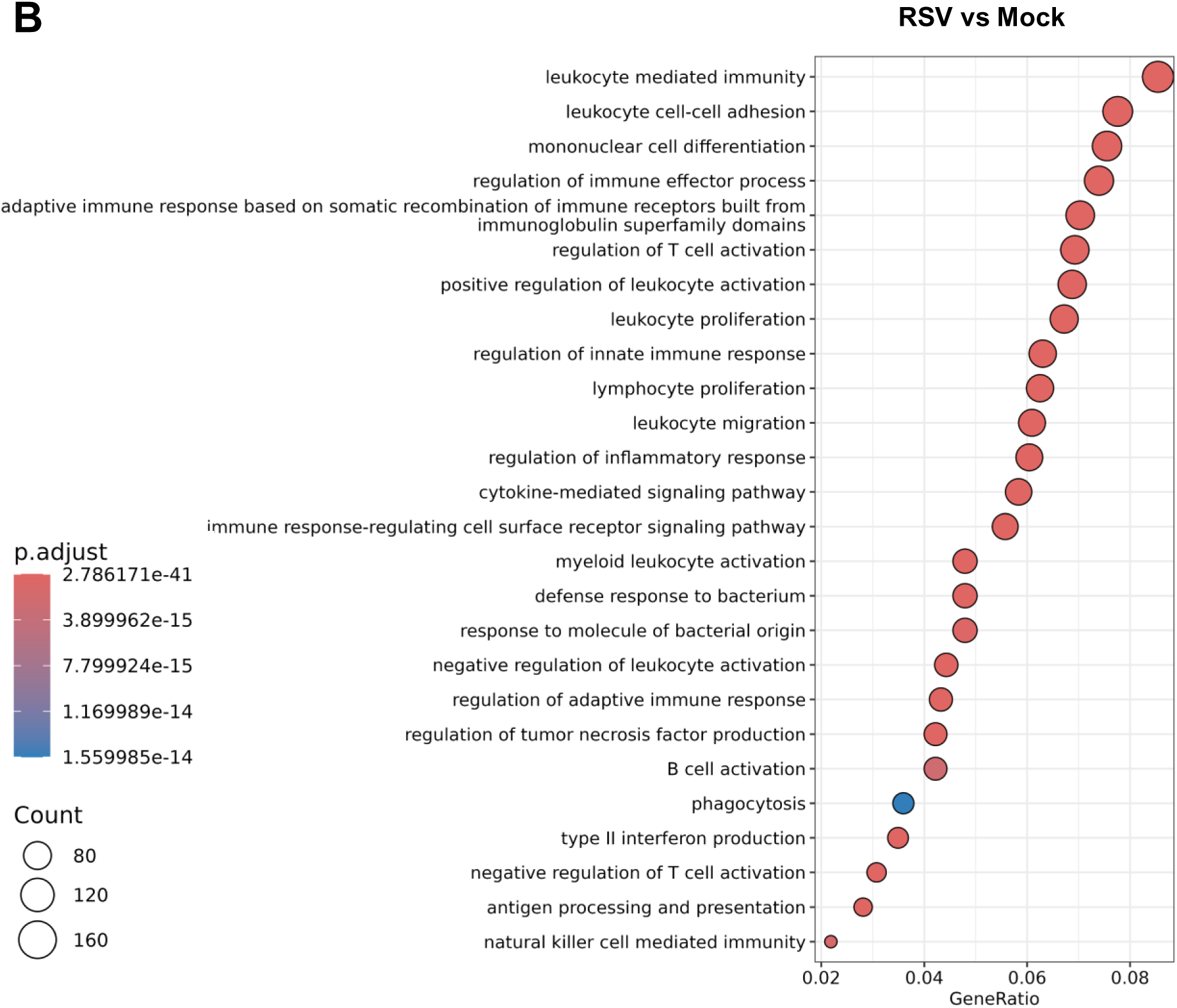
Transcriptome analysis of lung samples of RSV-infected mice. B6 mice were infected i.n. with RSV or mock, and on day 9 post infection, lung tissue RNA was isolated for RNAseq analysis. **A.** Heatmap of the top 100 differentially expressed genes in RSV-infected lung tissues compared to the mock controls at day 9. **B.** A simplified Gene Ontology (GO) enrichment dot plot showing the most significant signaling pathways induced by RSV infection compared to mock infection.

To validate RNAseq results, we next measured lung T cell responses in RSV-infected mice. Lung leukocytes were isolated on day 9 or day 30 post RSV infection. As shown in **Figure 5A-B**, ELISPOT assay showed that RSV induced more than 16-fold higher SARS-CoV-2-spike (S) protein specific T cell responses than the mock-infected mice. RSV-infected mice also displayed 3 and 8 - fold-higher CD69 (early T cell activation marker) expression on lung CD4^+^ and CD8^+^ T cells respectively compared to that of the mock-infected (**Figure 5C**). However, SARS-CoV-2-specific IgG and IgA antibodies in sera or bronchoalveolar lavage (BAL) fluid of RSV-infected BALB/c mice or B6 mice were barely detectable at day 9 (**Supplementary Figure 4A-D**). On day 30 post RSV infection, SARS-COV-2-reactive lung T cell responses were diminished and were 3-fold higher than the mock group (**Figure 5D**). CD69 expression on lung CD4^+^ and CD8^+^ T cells of RSV-infected mice was reduced and no significant differences were noted between those of RSV and mock-infected mice, suggesting a decreased T cell activation at the later time point (**Figure 5E**). No detectable levels of SARS-CoV-2 specific IgG antibodies were detected in sera of RSV-infected mice at day 30 (**Supplementary Figure 4E**). Thus, RSV induced SARS-CoV-2-reactive pulmonary T cell at day 9 but declined at day 30. As transcriptomic analysis indicates the induction of APCs and innate immune signaling pathway following RSV infection, to validate these results, we examined lung APC activation by measuring MHC class II expression on lung macrophages (F4/80^+^), dendritic cells (DCs, CD11c^+^), and epithelial cells (CD326^+^) in RSV-infected BALB/c mice. There was a 1.5-to-2.7-fold increase on levels of MHC II expression on lung macrophages, DCs, and epithelial cells at day 9 compared to the mock-infected mice (**Figure 6A**). The magnitude of MHC class II induction was reduced on DCs and lung epithelial cells at day 30 but remain unchanged on lung macrophages (**Figure 6B**). Furthermore, at day 9 post RSV infection, mice were challenged with SARS-CoV-2 mouse adaptive strain. On day 2, there was a 1.5-fold higher MHC class II expression on SARS-CoV-2 infected mice with RSV prior infection than those with mock prior infection, indicating a trained immunity induced by RSV infection (**Figure 6C**).

**Figure 5.**
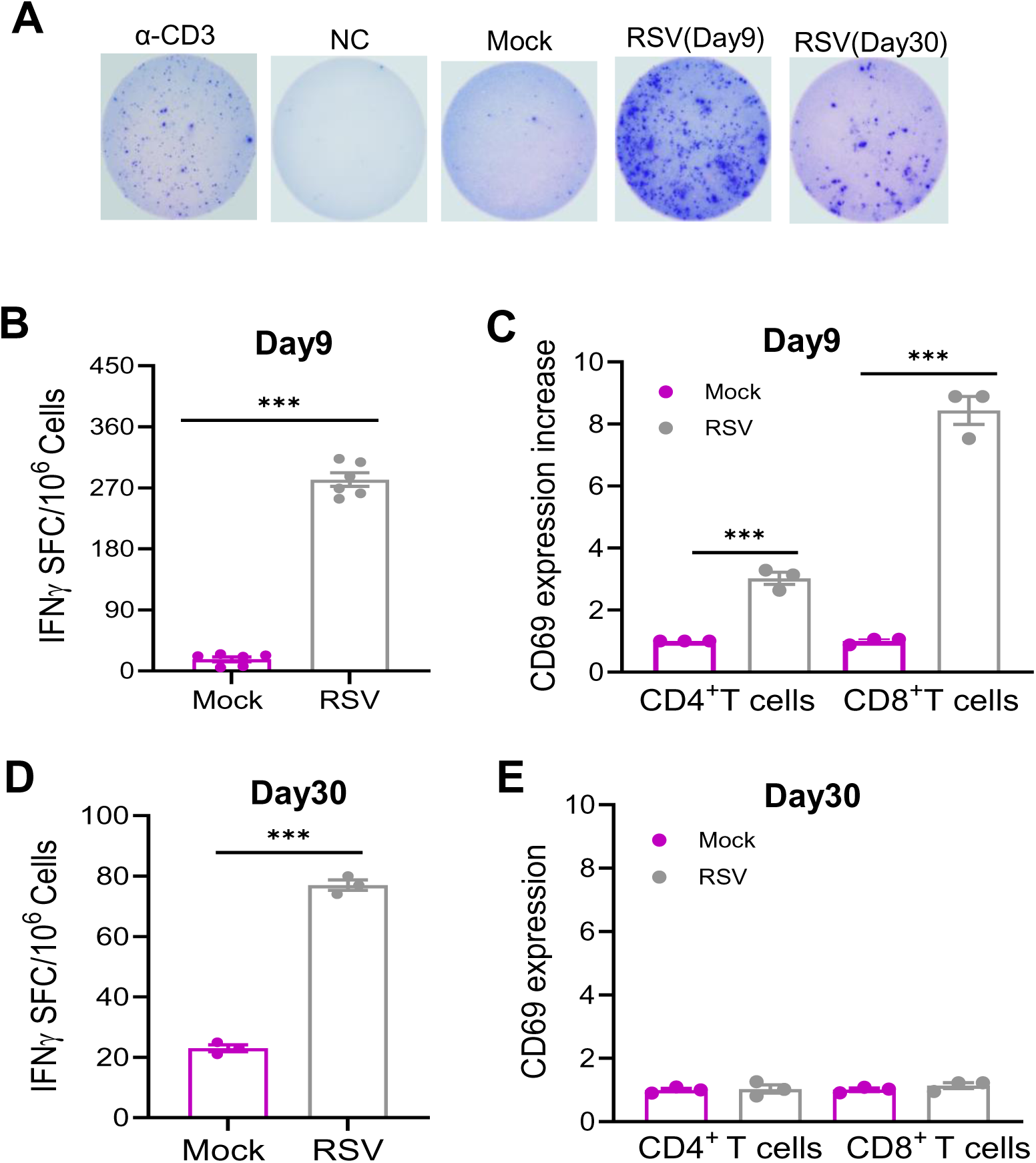
RSV induced SARS-CoV-2 reactive mucosal T cell responses. BALB/c mice were infected i.n. with RSV or PBS (mock), and on day 9 (**B-C**) or day 30 (**D-E**) post infection. Lung leukocytes were stimulated were stimulated with SARS-CoV-2 S & N peptides, α-CD3, or blank for 36 h. **A.** Images of wells from T cell culture. **B** & **D**. Spot forming cells (SFC) were measured by IFN-γ ELISpot. Data are shown as # of SFC per 10^6^ lung leukocytes. n= 3 to 6. **C** & **E**. CD69 expression on lung T cell subsets. Cells were gated in total lung leukocytes. Data are presented as fold increase compared to the mock-infected. \*\*\**P* < 0.001 RSV-infected compared to mock infected.

**Figure 6.**
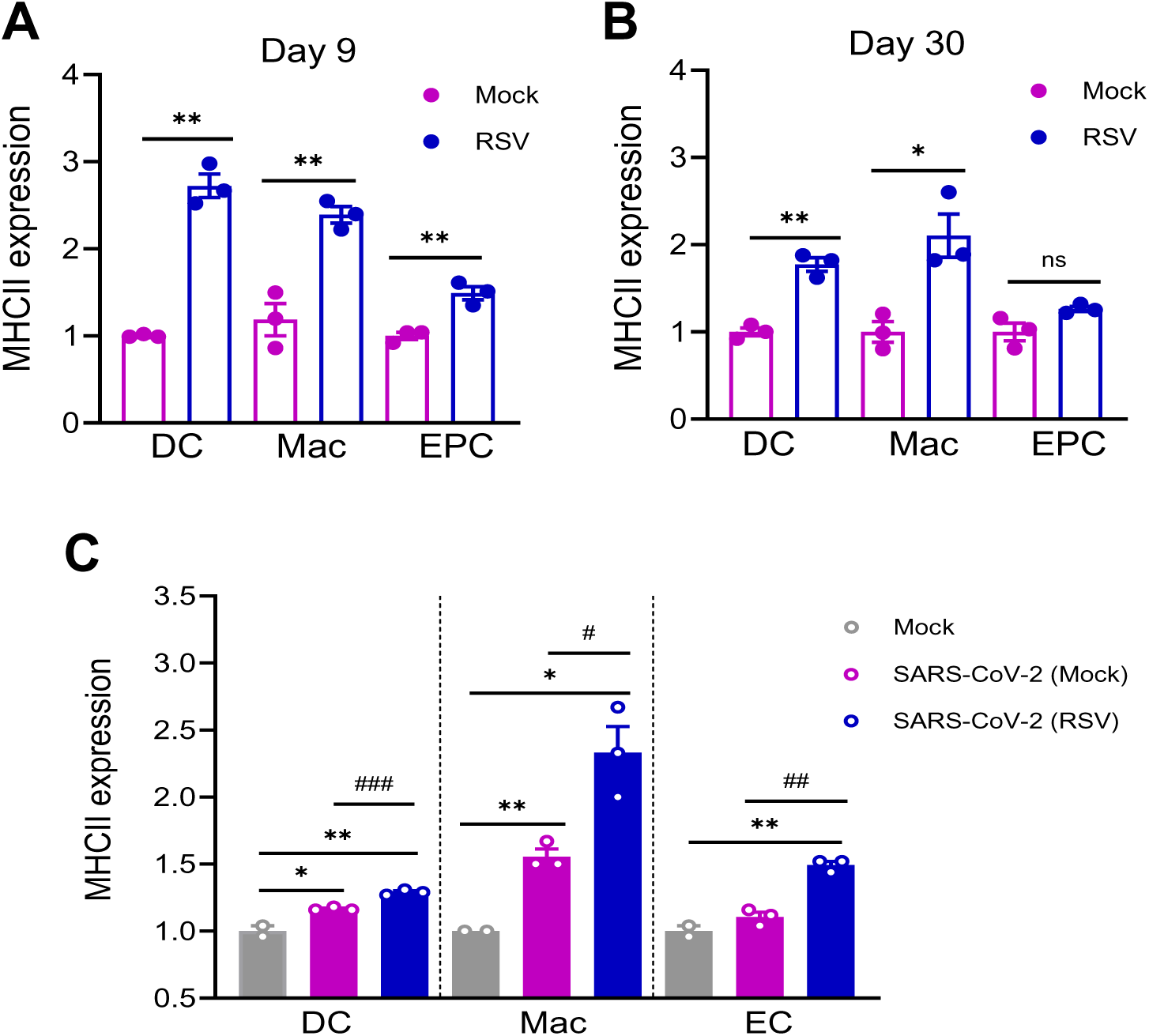
RSV-induced the activation of antigen presentation cells (APCs) in the lung. **A-B.** BALB/c mice were infected i.n. with RSV or PBS (mock). On day 9 (**A**) and day 30 (**B**) post infection, lung APCs were stained for CD11c (DC, dendritic cells), F4/80 [(macrophages),Mac] CD326 (epithelial cells, EC) and MHC class II. **C.** B6 mice were infected i.n. with RSV or mock, and on day 30 post infection, mice were challenged with mouse-adaptive SARS-CoV-2 strain CMA4. Lung leukocytes were isolated at day 2 post SARS-CoV-2 challenge and stained for APC markers (CD11C, F4/80, CD326) and MHC II. Cells were gated in total lung leukocytes. Data are presented as fold increase compared to mock-infected. n= 2 to 3. Data are presentive of two similar experiments. ** *P* < 0.01, or \**P* < 0.05 compared to mock group. ^###^*P* < 0.001, ^##^*P* < 0.01, or ^#^*P* < 0.05 compared to SARS-CoV-2 (mock) group.

### RSV induces heterologous protection against subsequent SARS-CoV-2 infection partially via modulation of γδ T cell functions

The cross talk between γδ T cells and APCs is known to play an important role in APC maturation. We noticed that lung γδ T cells expanded more than 6-fold by day 9 (**Figure 7A**). and reduced to 2-fold by day 30 in RSV-infected BALB/c mice, coinciding with the waning RSV-induced protection against SARS-CoV-2 infection. SARS-CoV-2 infection in B6 mice triggered 1.6-fold higher expansion of γδ T cells in mice-prior infected with RSV than those prior mock-infected (**Figure 7B**). These results suggested that a trained γδ T cell response triggered by RSV infection.

**Figure 7.**
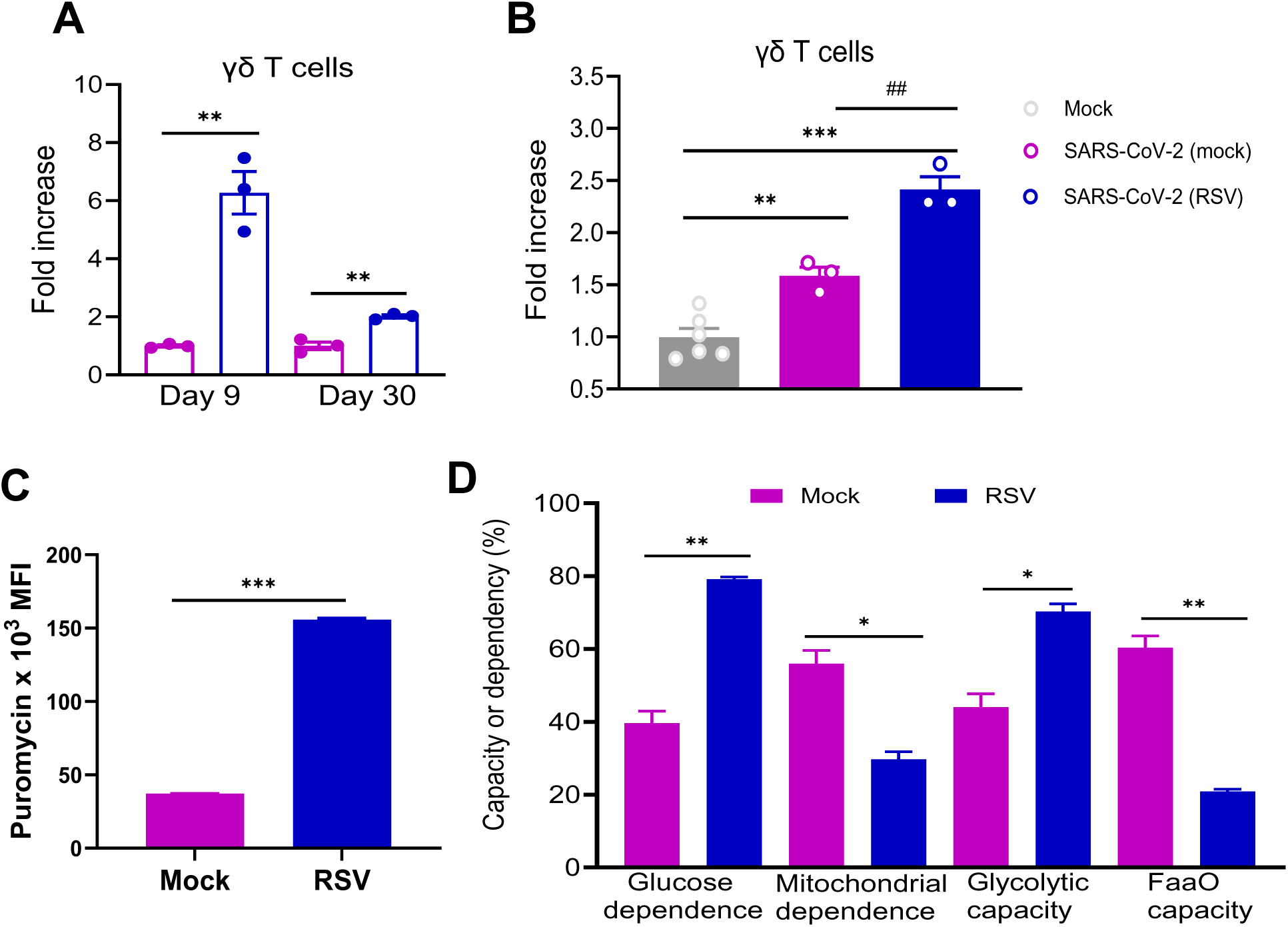
RSV induced trained γδ T cell responses. **A.** BALB/c mice were infected i.n. with RSV or PBS (mock). On day 9 and day 30 post infection, lung leukocytes were stained for CD3 and TCRγδ. Cells were gated in total lung leukocytes for percent positive and data are presented as fold increase compared to mock-infected. **B.** B6 mice were infected i.n. with RSV or mock, and on day 30 post infection, mice were challenged with mouse-adaptive SARS-CoV-2 strain CMA4. Lung leukocytes were isolated at day 2 post SARS-CoV-2 challenge and stained for CD3 and TCRγδ. Cells were gated in total lung leukocytes for percent positive and data are presented as fold increase compared to mock-infected. n =3 to 6. **C-D.** Lung leukocytes were isolated from RSV or mock-infected BALB/c mice at day 9 pi. Metabolic parameters by modified SCENITH (https://www.scenith.com), calculated as described in (24). (**C**) puromycin incorporation, (**D**) Glycolytic capacity, fatty acid oxidation/amino acid oxidation (FAO/AAO) capacity, mitochondrial dependence, and glucose dependence. All parameters were measured by flow cytometry. Data are one representative of three similar experiments. *** *P* < 0.001, ** *P* < 0.01, or \**P* < 0.05 compared to mock group. ^##^*P* < 0.01 compared to SARS-CoV-2 (mock) group.

We next studied the functional metabolism profile of lung γδ T cell in RSV-infected mice utilizing the SCENITH method ^19^, a flow cytometry–based technique using puromycin incorporation as a proxy for protein synthesis activity and ATP usage by the cell. RSV infection induced a substantial increase in protein synthesis levels (puromycin MFI, **Figure 7C**) on γδ T cells. We further calculated fatty acid/amino acid oxidation capacity, glycolytic capacity, mitochondrial dependence, and glucose dependence of lung γδ T cells by using the treatment with metabolic inhibitors and found that lung γδ T cells of RSV-infected mice had an increased glycolytic capacity with a decreased dependence on mitochondrial energy metabolism (**Figure 7D, Supplementary Figure 5A-B**). In addition, there was a decreased fatty acid/amino acid oxidation capacity with an increased glucose dependence in lung γδ T cells of RSV-infected mice. To determine whether γδ T cells contributed to induction SARS-CoV-2 reactive T cell responses, we infected WT B6 and TCRδ^-/-^ mice i.n. with 5 x10^6^ PFU RSV A2 or mock. At day 9, RSV-infected WT and TCRδ^-/-^ mice both showed induction of SARS-CoV-2-specific Th-1 prone immune responses in the lung compared to their respective mock-infected controls. There was more than 1-fold-higher SARSCoV-2-reactive T cell response in the lung of WT mice than those of TCRδ^-/-^ mice, suggesting RSV-induced SARS-CoV-2 specific pulmonary T cells partially dependent on γδ T cells (**Figure 8A**). There were barely any detectable SARS-CoV-2 specific IgA^+^ B cells induced in the lungs of either group (**Supplementary Figure 5C**). In the spleen, no significant differences were noted between WT and TCRδ^-/-^ mice, though a low SARS-CoV-2 specific T cell response was noted in both groups (**Figure 8B**). Lastly, to understand whether γδ T cells contribute to RSV-induced heterologous protection, we infected WT B6 mice and TCRδ^-/-^ mice i.n. with 5 x10^6^ PFU RSV A2 or mock. Mice were monitored daily for weight changes and morbidity. TCRδ^-/-^ mice showed more weight loss within 5 days post RSV infection than WT group (**Supplementary Figure 5D**). At day 9 of RSV infection, all mice were challenged with 1 × 10^4^ PFU mouse-adapted SARS-CoV-2 strain CMA4. WT mice with prior RSV infection had significantly lower levels of infectious particles and viral RNA than WT mice with prior mock infection (**Figure 8C-D**). Although TCRδ^-/-^ mice with RSV prior infection also showed reduced viral RNA levels compared to TCRδ^-/-^ mice with prior mock infection, there were significantly higher levels of infectious particles and viral RNA noted in TCRδ^-/-^ group compared to WT mice with prior RSV infection. Furthermore, TCRδ^-/-^ mice with RSV prior infection also had increased levels of chemokines than WT mice with prior RSV infection (**Figure 8E-G**), indicating more lung pathology in the absence of γδ T cells. Together, these results suggest that RSV reprogramed the metabolic profile of lung γδ T cells, which in turn contribute to the development of SARS-CoV-2 reactive T cells and heterologous protection against SARS-COV-2 challenge.

**Figure 8.**
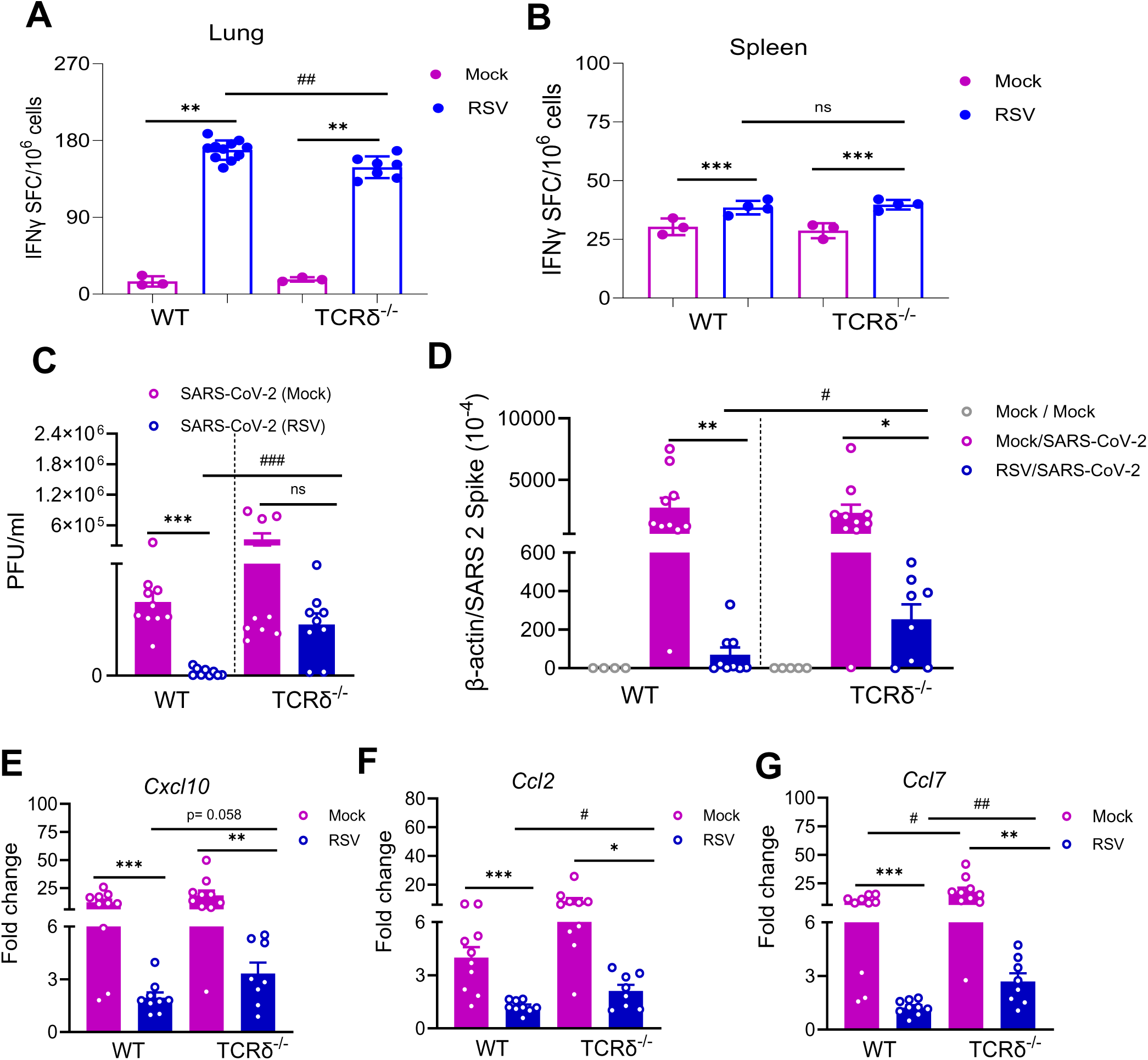
RSV-induced γδ T cell activation contributed to heterologous protection against subsequent SARS-CoV-2 challenge. WT B6 and TCRδ^-/-^ mice were infected i.n. with 5 x10^6^ PFU RSV A2 or mock. **A-B.** Lung leukocytes (**A**) or splenocytes (**B**) were stimulated with SARS-CoV-2 S & N peptides, or blank for 24 to 36 h. Spot forming cells (SFC) were measured by IFN-γ ELISpot. Data are shown as # of SFC per 10^6^ lung leukocytes. n= 3 to 11. **C-G**. At day 9 of RSV infection, all mice were challenged with 1 × 10^4^ PFU mouse-adapted SARS-CoV-2 strain CMA4. Two days after viral challenge, lung tissues were collected. SARS-CoV-2 viral titers in lung tissues were measured by plaque (**C**) and Q-PCR (**D**) assays. **E-G**. Measurement of chemokine levels in lung tissues by Q-PCR assays at day 2 post infection. Data are presented as the fold increase compared to naïve mice (means ± SEM). n= 8 to 10. *** *P* < 0.001, ** *P* < 0.01, or \**P* < 0.05 compared to mock group. ^###^*P* < 0.001, ^##^*P* < 0.01, or ^#^*P* < 0.05 compared to WT group.

## DISCUSSION

In this study, we demonstrated in two animal models that prior RSV infection induces heterologous protection against subsequent SARS-CoV-2 infection challenge via induction of γδ T cell-mediated trained immunity. The heterologous protection is dose and time-dependent. Trained immunity is also called innate immune memory. Several innate immune cells, including monocytes, macrophages, neutrophils, DCs, and natural killer cells, have been reported to be involved in trained immunity ^4,5^. gd T cells were recently reported to play a key role in trained immunity induced by MMR vaccination ^20^. Here, we found that RSV infection triggered γδ T cell expansion and modified its cellular metabolic pathway.

γδ T cells represent a minority of the CD3^+^ T cells in lymphoid tissue and blood but are well enriched at epithelial and mucosal sites. These cells lack major histocompatibility complex (MHC) restriction but express innate-like receptors, such as toll-like receptor (TLR) and NKG2D features, play an important role in innate immunity ^21^. Clinical and animal studies suggest that γδ T cells are involved in host protection against SARS-CoV-2 infection ^22–24^. Here, we found that TCRδ^-/-^ mice showed more weight loss within 5 days post RSV infection than RSV-infected WT mice. In addition, TCRδ^-/-^ mice with RSV prior infection also had increased levels of chemokines than WT mice with prior RSV infection. These data suggest that γδ T cells contribute to protective immunity against RSV. More importantly, RSV-trained γδ T cells also contribute to host protection against SARS-CoV-2 challenge. γδ T cells are also known to cross-talk with APCs and further promote their maturation and T cell priming and enhance memory T cell development during microbial infection ^25–27^. We also noted trained lung APC responses, including DCs, macrophages, and epithelial cells. Furthermore, RSV-infected TCRδ^-/-^ mice showed reduced SARS-CoV-2 reactive T cells. Together these results suggest that RSV-induced γδ T cells contribute to lung APC activation and further T cell priming.

gd T cells are also known to mediate humoral immunity and facilitate CD19^+^ B cell activation and the production of immunoglobulins ^28,29^. Furthermore, γδ T cell expansion was also associated with higher anti-SARS-CoV immunoglobulin G titers ^30^. We found that RSV infection induced minimal levels of SARS-CoV-2 reactive B cells and low antibody responses in the periphery and mucosal sites. It’s likely that RSV-induced γδ T cell expansion does not result in SARS-CoV-2-specific antibody responses. Furthermore, TCRδ^-/-^ mice with RSV prior infection displayed reduced but not abolished heterologous protection from SARS-CoV-2 infection and SARS-CoV-2 reactive mucosal T cells. Thus, γδ T cell-independent, likely RSV-induced trained Lung APCs directly contribute to the induction of SARS-CoV-2 reactive T cells, leading to protection of the host from SARS-CoV-2 challenge.

Two animal models, including BALB/c mice and K18-hACE2 mice were used to study RSV-induced heterologous protection against SARS-CoV-2. As K18 hACE2 mice were generated in B6 background, further immunological analysis was performed in both BALB/c mice and C57BL6 mice. We demonstrated that RSV-induced SARS-CoV-2 reactive T cells in both mouse models and declined over time. Furthermore, RSV-induced trained immunity, including lung APCs and γδ T cells also declined at 1 month. These data suggest RSV-induced trained immunity is transient and associated with its heterologous protection. Using both animal models allows us to assess how the virus behaves in a natural mouse-adapted environment versus a humanized system. Furthermore, these studies offer a broader understanding of RSV-induced trained immunity due to their distinct genetic and immunological backgrounds.

Prior vaccination or viral infection with other respiratory viruses can influence the trajectory of SARS-CoV-2. Influenza and BCG vaccines were reported to protect SARS-CoV-2 infection via induction of trained immunity ^31–34^. However, the impact of sequential infection with other viruses (e.g., RSV, Influenza, and hMPV) on vulnerability to SARS-CoV-2 disease or vice versa remains unknown. Results from this study will advance our understanding of the pathogenesis and host immunity of the co-circulation of other respiratory viruses with SARS-CoV-2 and its variants. Trained immunity can be used as a strategy to boost vaccine efficacy and enhance host specific and non-specific immune responses. Investigating the impact of recent infection on SARS-CoV-2 infection and vaccination efficacy will provide new insights into SARS-CoV-2 pathogenesis as well as future vaccine strategies.

## METHODS

### Viruses

SARS-CoV-2 USA-WA1/2020 strain was obtained from the World Reference Center for Emerging Viruses and Arboviruses (WRCEVA) at the University of Texas Medical Branch (UTMB) and was amplified twice in Vero E6 cells. Mouse-adapted SARS-CoV-2 strain CMA4 viral stocks were provided by Dr. Xuping Xie at UTMB. The generation of SARS-CoV-2 strain CMA4 was described previously ^14^. RSV long strain was grown in HEp-2 cells and purified by sucrose gradient as described ^35^.

### Mice infection and tissue collection

7-to 8-week-old BALB/c and C57BL/(B)6 mice were purchased from Jackson Laboratory. γδ T cell-deficient mice (TCRδ ^-^/^-^) and K18 hACE2 mice (stock #034860, Jackson Laboratory) were both on B6 background and were bred at the UTMB animal facility. Age- and sex-matched male and female mice were used in this study. Mice were infected intranasally (i.n) with 5 × 10^6^ or 1 × 10^7^ plaque-forming units (PFU) of RSV. At day 9 or day 30 post RSV infection, mice were challenged with 1 × 10^4^ PFU of the SARS-CoV-2 CMA4 strain or 2 × 10^3^ PFU of the SARS-CoV-2 USA-WA1/2020 strain. Infected mice were monitored twice daily for morbidity and mortality. In some experiments, infected mice were euthanized for tissue collection. The right superior and inferior lobes of lung tissues were then collected in Trizol for RNA extraction and in DMEM for viral titration by plaque assay, respectively ^14^. All animal experiments were approved by the Animal Care and Use Committees at UTMB.

### Quantitative PCR (Q-PCR)

Lung tissues were resuspended in TRIzol for RNA extraction according to the manufacturer’s instructions (Thermo Fisher). Complementary (c) DNA was synthesized by using a qScript cDNA synthesis kit (Bio-Rad, Hercules, CA). The sequences of the Q-PCR primer sets for mouse cytokines, chemokines, SARS-CoV-2 S gene and PCR reaction conditions were described previously ^36–38^. The PCR assay was performed in the CFX96 real-time PCR system (Bio-Rad). Gene expression was calculated using the formula 2^ ^-[C^t^(target^ ^gene)-C^t^(*β-actin*)]^ as described before ^39^.

### Plaque assay

Vero E6 cells were seeded in 6-well plates and incubated at 37°C. 10-fold serially diluted lung tissue homogenates were used to infect cells at 37°C for 1 h. After the incubation, cells were overlaid with 2X-MEM (Thermo Fisher) with 8% FBS and 1.6% agarose (VWR). After 48 h incubation, plates were stained with 0.05% neutral red (Sigma-Aldrich) and plaques were counted to calculate virus titers expressed as PFU/ml.

### Antibody ELISA

ELISA plates (Corning, USA) were coated with 100 ng/well recombinant SARS-CoV-2 RBD protein (RayBiotech) for overnight at 4°C. The plates were washed twice with PBS containing 0.05% Tween-20 (PBS-T) and then blocked with 8% FBS for 1.5 h. Sera were diluted 1:100 in blocking buffer. Bronchoalveolar lavage (BAL) fluid was collected by making an incision in the trachea and lavaging the lung twice with 1 mL PBS. Following centrifugation, supernatant was collected to measure antibody titers. BAL fluid was diluted 1:3 in blocking buffer. Samples were added for 1 h at 37°C. Plates were washed 5 times with PBS-T. Goat anti-mouse IgG (Southern Biotech) coupled to horseradish peroxidase (HRP) or alkaline phosphatase, For IgA measurement, goat anti-mouse IgA (Southern Biotech) coupled to HRP was added at a 1:2000 dilutions for 1 h at 37°C. The was followed by adding TMB (3, 3, 5, 5′-tetramethylbenzidine) peroxidase substrate (Thermo Fisher Scientific) for about 5 min. The reactions were stopped by 1M sulfuric acid, and the intensity was read at an absorbance of 450nm.

### Flow cytometry

Lung leukocytes were stained with antibodies for CD3, TCRγδ, CD4, CD8, CD69, F4/80, CD11c, CD326, and MHC II (all antibodies were purchased from ThermoFisher). After staining, the cells were fixed in 1% paraformaldehyde and acquired by a C6 flow cytometer instrument (BD Biosciences). Dead cells were excluded based on forward and side light scatter. Data were analyzed with a CFlow Plus flow cytometer (BD Biosciences).

### IFN-γ ELISPOT

Millipore ELISPOT plates (Millipore Ltd) were coated with mouse anti-IFN-γ capture Ab at 1:100 dilution (Cellular Technology Ltd) and incubated at 4°C overnight. Lung leukocytes or splenocytes were stimulated with SARS-CoV-2 S and nucleocapsid (N) peptide pools (2 μg/ml, Miltenyi Biotec) for 36 h at 37°C. Cells were stimulated with anti-CD3 (1 μg/ml, e-Biosciences) or medium alone were used as positive and negative controls. This was followed by incubation with biotin-conjugated anti-IFN-γ at 1:100 dilution (Cellular Technology Ltd,) for 2 h at room temperature, followed by incubation with alkaline phosphatase-conjugated streptavidin for 30 min. The plates were washed and scanned using an ImmunoSpot 6.0 analyzer and analyzed by ImmunoSpot software to determine the spot-forming cells (SFC) per 10^6^ lung leukocytes.

### Energy metabolism profile of lung gd T cells

Metabolism profile analysis was performed as described previously with some modifications ^19,20^. Briefly, 1 x 10^6^ lung leucocytes of mock or RSV-infected mice were stimulated with PMA (Sigma, 50 ng/ml) and Ionomycin (Sigma, 500 ng/ml) for 4 h. Cells were then either left untreated (control, Co) or treated with 2-deoxy-D-glucose (DG) (100 mM), oligomycin (O) (10 μM), and a combination of 2-DG and oligomycin (DGO) (100 mM and 10 μM) for 30 min. Following the addition of puromycin (10 μg/mL), the cells were incubated for an additional 45 min, and the cells were subsequently harvested and washed in cold FACS buffer before being stained with antibodies for TCRgd and CD3 (ThermoFisher) for 20 min in ice. Cells were washed, fixed in 2% paraformaldehyde, and permeabilized with 0.5% saponin before adding mouse anti-puromycin-PE antibody (BioLegend, USA). Samples were acquired by a C6 Flow Cytometer instrument. Dead cells were excluded based on forward and side light scatter. Data were analyzed with a CFlow Plus Flow Cytometer (BD Biosciences). Based on puromycin mean fluorescence intensity (MFI) on gated gd T cells, several parameters of the energetic metabolism profile metabolic, including glucose dependence, mitochondrial dependence, glycolytic capacity, and fatty acid and amino acid oxidation (FaaO) capacity were calculated as described previously ^19^.

### RNA-seq analysis

RNA was extracted from lung tissues as described above and 2 µg of RNA was used for RNA-seq analysis. RNA samples quality was assessed using Agilent Bioanalyzer RNA Nano Chips (Agilent, Santa Clara, CA). RNAseq libraries were then prepared using NEBNext rRNA depletion kit v2 (Cat# 7400) and Ultra II Directional RNA library preparation kit (Cat# 7760) (NEB, Ipswich, MA) following manufacturer’s recommended procedure. The resulting libraries were run on Agilent Bioanalyzer High Sensitivity DNA Chips (Agilent, Santa Clara, CA) for size and quantified using qPCR. Sequencing was carried out on Element Biosciences Aviti sequencer (Element Biosciences, San Diego, CA) using paired end 75bp parameter to a sequencing depth of above 50 million paired reads per sample (57 million to 93 million). The RNA-seq data in this study were deposited in NCBI’s Gene Expression Omnibus (GEO Series accession number GSE300910). The reads were quality filtered and trimmed for adapter sequence using Trimmomatic-0.39 ^40^ and aligned to mouse GRCm39 reference genome using STAR 2.7.11a ^41^. Differential expression was performed using Bioconductor DESeq2 package ^42^ and GO enrichment analysis were performed using Bioconductor clusterProfiler package ^43^. GSEA analysis was performed using GSEA version 3.0 ^44^.

### Statistical analysis

Survival curve comparison was performed using GraphPad Prism software 9.4.1, which uses the log-rank test. Values for weight changes, viral load, cytokine production, antibody titers, and T cell response experiments were compared using Prism software statistical analysis and were presented as means ± SEM. P values of these experiments were calculated with a non-paired Student’s t test.

## ACKNOWLEDGEMENTS

This study was supported in part by National Institute of Health grants R01AI127744 (TW), R01 NS125778 (TW), R01 AI176670 (TW), R21 AI178135 (TW), ERP-1252718 ALA (XB), R21 AI166543 (XB), R61 AG075725 (XB), and a pilot grant supported by the Institute for Human Infections and Immunity at UTMB (TW & XB),

## AUTHOR CONTRIBUTIONS

AA, WW, and MJ performed the experiments. AA. WW, XB, and TW designed the experiment. AA, MJ, HH, A, PS, and TW analyzed the data. AA, HH, A, and TW wrote the first draft of the manuscript, and the other authors edited the manuscript.

## SUPPLEMENTARY FIGURE LEGENDS

**Supplementary Figure 1: Weight loss after RSV infection in K18-hACE2 mice.** K18-hACE2 mice were infected with a low (LD) or high dose (HD) of RSV or mock. Mice were monitored daily for weight loss. Weight loss is indicated by percentage using the weight on the day of infection as 100%. \*\*\**P* < 0.001, \*\**P* < 0.01, or \**P* < 0.05 LD (n= 23) or HD (n =10) group compared to mock-infected mice (n = 18).

**Supplementary Figure 2: RNAseq analysis of lung samples of RSV-infected mice.** B6 mice were infected i.n. with RSV or mock. Lung tissues were collected at day 9 post RSV or mock infection. Lung tissue RNA was used for RNAseq analysis. **A.** Volcano plot of RSV vs. Mock groups. Color dots represent genes that meet different criteria of analysis. Gray color: No significance (does not meet either significance *P* value or foldchange cut off); Green dots: Meeting only foldchange cutoff; Blue dots: meeting *P* value cutoff (multi-testing adjusted *P* < 0.1); Red dots: meeting both fold change and *P* value cutoff. **B-D.** GSEA enrichment plots for immune-related pathways in RSV vs. mock group. Enrichment plots show upregulation of (**B**) lymphocyte-mediated immunity, (**C**) leukocyte-mediated immunity, and adaptive immune response pathways in RSV-infected samples compared to mock. Normalized enrichment scores (NES) and false discovery rates (FDR) are indicated for each pathway. Heatmaps depict the relative expression levels of genes within (**B**) lymphocyte-mediated immunity, (**C**) leukocyte-mediated immunity, and (**D**) adaptive immune response pathways. Each row represents a gene, and each column represents a sample. Red indicates higher expression; blue indicates lower expression.

**Supplementary Figure 3: RNAseq analysis of lung samples of SARS-CoV-2-infected mice with prior RSV infection.** B6 mice were infected i.n. with RSV or mock, and on day 9 post infection, mice were challenged with SARS-CoV-2 CMA4. Lung tissues were collected at day 2 post SARS-CoV-2 challenge in mice with prior RSV or mock infection. Lung tissue RNA was used for RNAseq analysis. A simplified GO enrichment dot plot showing the most significant signaling pathways induced by SARS-CoV-2 challenge with prior RSV infection compared to SARS-CoV-2 infected mice with prior mock infection.

**Supplementary Figure 4: SARS-CoV-2 specific antibody responses in RSV-infected mice.** BALB/c (**A-B** & **E**) or B6 (**C-D**) mice were infected i.n. with RSV or mock. At day 9 (**A-D**) or day 30 (D) post infection, sera and BAL were collected. Results of IgG and IgA ELISA using SARS-CoV-2 recombinant RBD protein antigen were presented.

**Supplementary Figure 5: RSV-induced γδ T cell responses contribute to heterologous protection against subsequent SARS-CoV-2 challenge. A-B.** Metabolic analysis by a modified SCENITH. Lung leukocytes were isolated from mock **(A)** or RSV **(B)**infected BALB/c mice at day 9 pi. Cells were gated on CD3^+^TCRγδ^+^ cells for puromycin incorporation, Data are presentive of three similar experiments. **C.** SARS-CoV-2 specific memory B cell (MBC) responses by ELISPOT analysis at day 9 post RSV infection. Frequencies of RBD specific ASCs per 10^6^ input cells in MBC cultures from the subject. **D.** WT B6 and TCRδ^-/-^ mice were infected i.n. with 5 x10^6^ PFU RSV A2 or mock. Mice were monitored daily for weight loss. Weight loss is indicated by percentage using the weight on the day of infection as 100%. \*\*\**P* < 0.001, \*\**P* < 0.01, or \**P* < 0.05 WT mice RSV (n= 23) or TCRδ^-/-^ mice RSV (n =19) groups compared to WT mice mock (n =16) or TCRδ^-/-^ mice mock groups (n= 13) respectively. ^##^*P* < 0.01, or ^#^*P* < 0.05, WT mice RSV (n= 23) compared to TCRδ^-/-^ mice RSV (n= 19).

## Supplementary Methods

### B cell ELISPOT

Millipore ELISPOT plates (Millipore Ltd, Darmstadt, Germany) were coated with 100 µl of 15 µg/ml rSARS-CoV-2 RBD protein (RayBiotech). To detect total IgA^+^-expressing B cells, the wells were coated with 100 µl of anti-mouse IgA capture Ab 15 µg/ml, (Mabtech In). Cells were added in duplicate wells to assess total IgA ASCs or SARS-CoV-2 specific B cells. The plates were incubated overnight at 37°C, followed by incubation with biotin-conjugated anti-mouse IgA (0.5 µg/ml, 3865-6, Mabtech In) for 2 h at room temperature, then 100 µL/well streptavidin-ALP (1:1000) was added for 1 h. Plates were developed with BCIP/NBT-Plus substrate until distinct spots emerge, washed with tap water, and scanned using an ImmunoSpot 6.0 analyzer and analyzed by ImmunoSpot software (Cellular Technology Ltd).

