## Supplementary Figures for "Respiratory syncytial virus infection confers heterologous protection against SARS-CoV-2 via induction of γδ T cell-mediated trained immunity and SARS-CoV-2 reactive mucosal T cells"

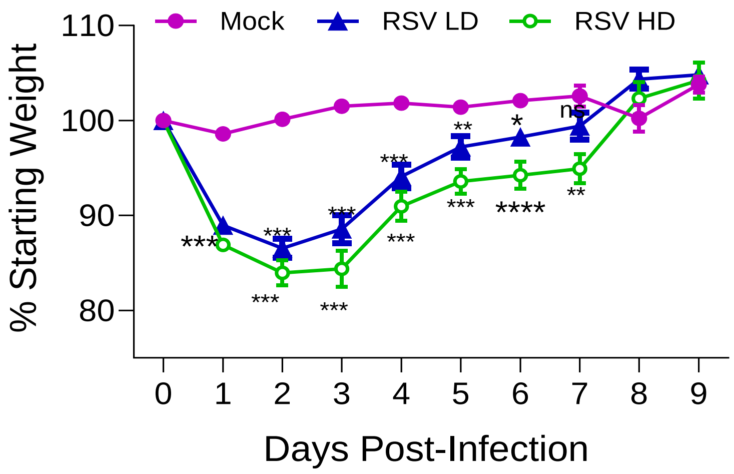


Supplementary Figure 1

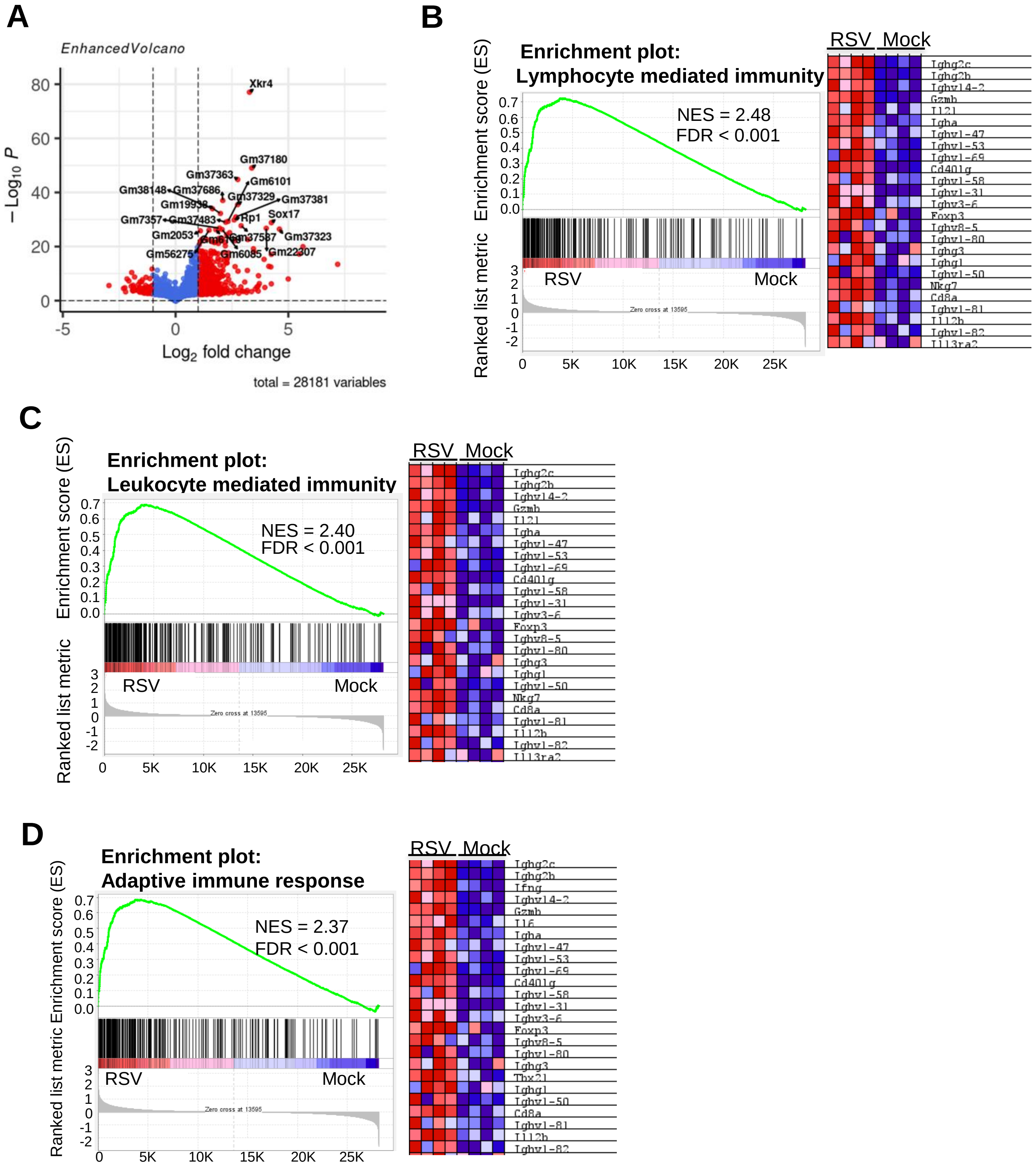


**Supplementary Figure 2**

### SARS-CoV-2 (RSV) Vs SARS-CoV-2 (Mock)

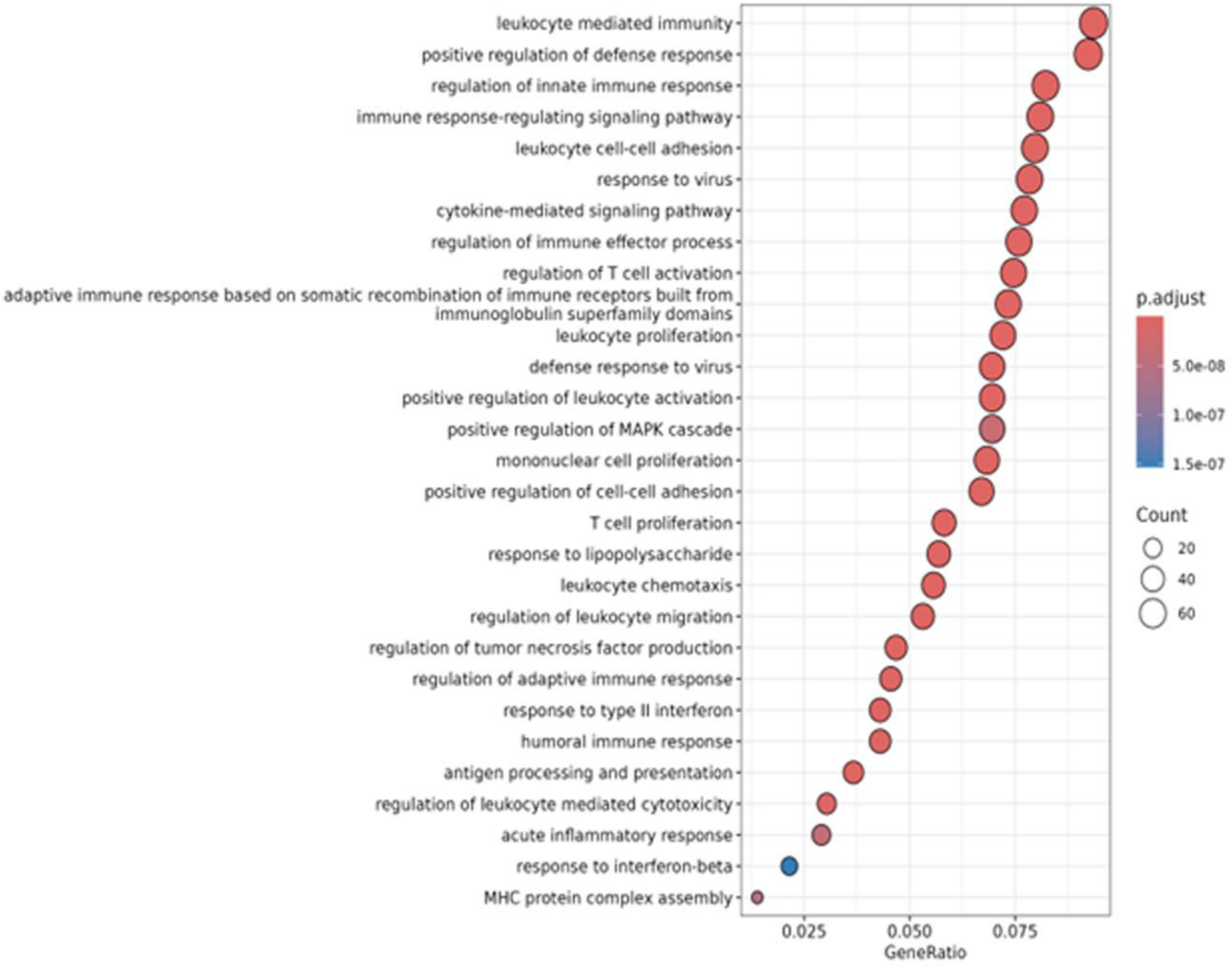


**Supplementary Figure 3**

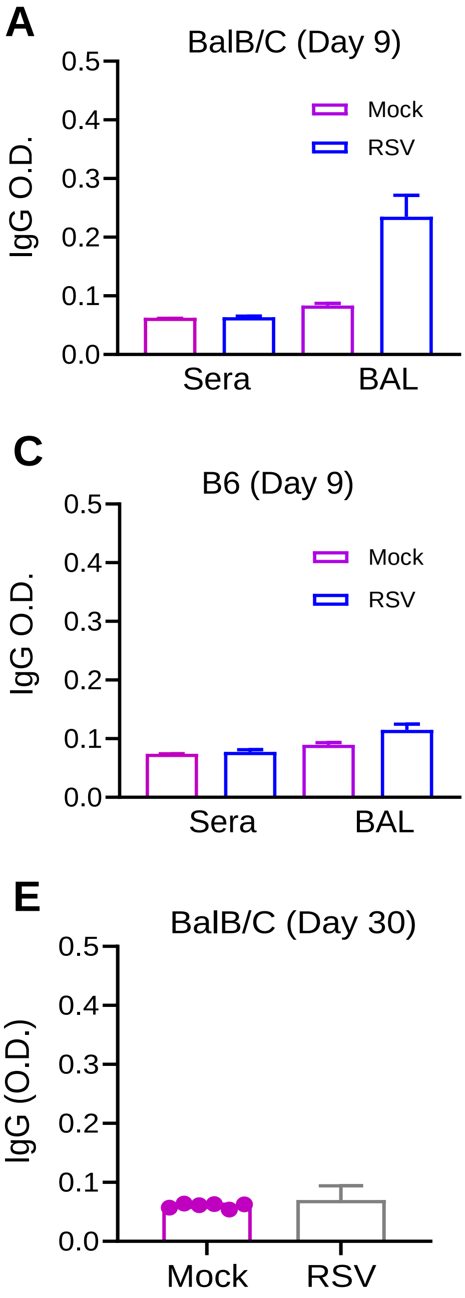


**Supplementary Figure 4**

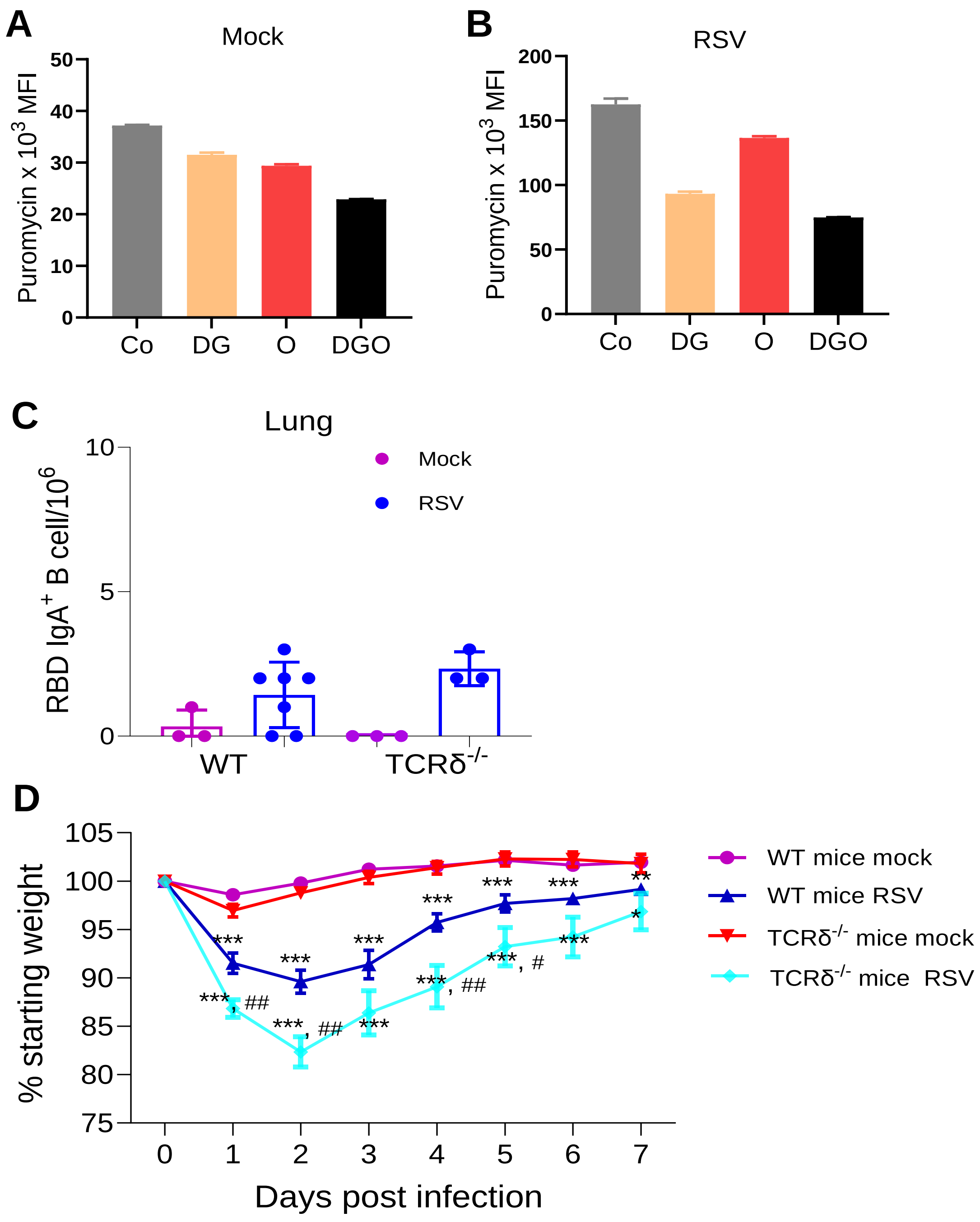


**Supplementary Figure 5**
